# Lake floodplains as sinks for stable soil organic carbon: is the carbon plant- or microbe-derived, and why does permanent land use matter?

**DOI:** 10.1101/2025.10.16.682620

**Authors:** Toky Jeriniaina Rabearison, Jim Felix-Faure, François Guillemette, Michelle Garneau, Christine Martineau, Louis Astorg, Martin Chantigny, Virginie Favreau, Jessika Malko, Alexandre Roy, Vincent Maire

## Abstract

Floodplain ecosystems play a key role in soil organic carbon (SOC) storage, as they integrate inputs from both vegetation and sediments. Promoting land uses with low anthropogenic disturbances helps maintain the function of these ecosystems as soil carbon (C) sinks. However, the tipping points along a disturbance gradient where land use transitions generate the largest SOC losses or gains remain unclear, though they are key for effective land use management and climate change mitigation. Because flood events can negatively impact labile C, determining the main origin of stable SOC, whether from plants or soil microorganisms, is also important to identify the optimal combination of land use, vegetation, and soil type for SOC stabilization in floodplains. We examined how SOC storage and stabilization vary in the floodplain of Lake Saint-Pierre, Quebec, Canada along an anthropogenic disturbance gradient of six land uses: conventional and improved croplands, temporary and permanent meadows, marshes, and forested swamps. In all 6 land uses, we quantified SOC stocks, mineral-associated organic matter - C (MAOM-C), particulate organic matter - C (POM-C), soil δ^13^C and sugars. Land use effects on SOC storage and stabilization were most pronounced in the topsoil (0-10 cm), with forested swamps showing the highest vegetation biomass, SOC, and MAOM-C. Microbe-derived inputs were more abundant in MAOM, whereas plant-derived C dominated POM. SOC gains increased with decreasing disturbance, with a tipping point occurring in the transition from temporary to permanent meadows. Our results highlight the importance of conserving permanent and low-disturbance land uses to promote SOC persistence in floodplain ecosystems and emphasize the central role of microbial metabolism in stabilization processes.

## 1 Introduction

Floodplain ecosystems are hotspots for soil organic carbon (SOC) storage due to their unique position at the interface between terrestrial and aquatic environments and their dynamic hydrological regimes (Mitra et al., 2005; Bridgham et al., 2006; Mitsch et al., 2013). Despite occupying only a small fraction of Earth’s surface, floodplain wetland soils play a disproportionately large role in global carbon (C) cycling by integrating both autochthonous organic inputs from vegetation and allochthonous materials deposited by sediment-rich floodwaters (Omengo et al., 2016; Scott and Wohl, 2018). These attributes render floodplains critical yet understudied components of the terrestrial C sink.

Although floodplains naturally emit greenhouse gas such as methane (Bridgham et al., 2006; Mitsch et al., 2013), their capacity to sequester soil C generally surpasses these emissions (Mitsch et al., 2013; Mitsch and Mander, 2018; Villa and Bernal, 2018). Maintaining or enhancing this C sink as well as transforming it into SOC stocks depends on choosing appropriate land uses (Ji et al., 2020). While intensive agriculture, such as annual cropping systems, can be particularly productive in floodplains (Kwesiga et al., 2019), repeated soil disturbance and harvest biomass exportation are conducive to reduced SOC stocks (Bronick and Lal, 2005; Ciais et al., 2010). In contrast, the conversion of croplands to forested floodplains has enhanced SOC due to higher litter inputs (Nyssen et al., 2008; Omengo et al., 2016; D’Elia et al., 2017) and increased transfers of SOC in deep soil layers through high fine root density (Ferré et al., 2014; Sariyildiz et al., 2022). However, the effect of forests on SOC stock compared to grasslands remains unclear in floodplain ecosystems, with some studies reporting higher SOC stock under forests (Smith and Reid, 2013; Hernandez et al., 2015), while others found higher SOC in grasslands due to better root development (Del Galdo et al., 2003; Sariyildiz et al., 2022) or reported no differences (Cierjacks et al., 2010; Scott and Wohl, 2018). These inconsistencies point to the need for a more nuanced understanding of land use effects along a gradient of anthropogenic disturbance in floodplain ecosystems.

Importantly, SOC responses to changes in land use are unlikely to be linear along the anthropogenic disturbance gradient. Tipping points may occur, where shifts in vegetation or management type can result in disproportionately large gains or losses in SOC storage (Lininger and Polvi, 2020; Yeasmin et al., 2020). For example, a transition from croplands to meadows may represent a stage where SOC gains are most pronounced, due to the cessation of tillage and crop residue removal (Ciais et al., 2010). Shifts in vegetation type and input quality, from herbaceous-dominated meadows to forests dominated by trees, could also lead to substantial changes in SOC storage (Filley et al., 2008; Córdova et al., 2018). Identifying the critical SOC gain along the disturbance gradient is essential to prioritize land use transitions that maximize SOC persistence in floodplains and support climate change mitigation.

Floodplains are also characterized by episodic flooding that can mobilize labile (.e., easily decomposed) forms of organic C (Saint-Laurent et al., 2017). As a result, the long-term stabilization of SOC is likely to depend on mineral-associated organic matter (MAOM), which forms through the adsorption of microbial by-products or plant-derived compounds onto mineral surfaces (Balesdent, 1996; Six et al., 2002; Cotrufo et al., 2013; Angst et al., 2021). MAOM is typically considered to be more persistent than particulate organic matter (POM), which is derived from partially decomposed plant materials (Cotrufo et al., 2013). MAOM accumulation is promoted in fine-textured soils, abundant in floodplain environments (Ricker and Lockaby, 2015; Heger et al., 2024). Nevertheless, its accumulation is not infinite and recent work by Georgiou et al. (2025) underlines the importance of considering MAOM saturation, i.e. its maximum observed capacity to stabilize C based on the physicochemical properties of the soil mineral matrix. Determining the degree of MAOM saturation in floodplains could therefore help to better recognize the potential of these ecosystems for storing more stable SOC.

Because plant- and microbe-derived compounds differ in their binding strength to minerals and vulnerability to decomposition, determining their proportion in MAOM composition is key to understanding and managing SOC persistence in floodplains (Kleber et al., 2021; Zhao et al., 2025). Yet, the main origin and composition of MAOM remain debated (Blankinship et al., 2018; Angst et al., 2021). Stable δ¹³C isotope can distinguish recent vegetation inputs from older carbon source (Balesdent and Mariotti, 1996; Tonucci et al., 2017), while sugar biomarkers help identify plant-vs microbe-derived compounds (Angst et al., 2021). Some studies have suggested predominant microbial residues in the MAOM (Cotrufo et al., 2013; Liang et al., 2017), whereas others have observed substantial plant-derived inputs (Angst et al., 2017; Córdova et al., 2018). Grasslands may favor microbial derived-amino sugars in MAOM compared to forests (Turrión et al., 2002) and croplands (Ding et al., 2017) likely due to continuous inputs of easily decomposed organic materials (Cotrufo et al., 2013; Angst et al., 2021). However, plant-derived neutral sugars in MAOM can be higher in forests than in croplands, but lower than in grasslands (Glaser et al., 2000; Jolivet et al., 2006). Understanding these patterns could facilitate the choice of land uses and vegetation types that promote long-term SOC stabilization in floodplain soils (Angst et al., 2021).

In this study, we investigated how SOC storage and stabilization vary across a floodplain under different land uses, along a disturbance gradient. Specifically, we aimed to trace the origin of MAOM in floodplain soils and to identify which land use transitions most effectively increase SOC and MAOM. We hypothesized that:

1. perennial ecosystems with low disturbance (forests, marshes and meadows) would outperform croplands in biomass inputs, SOC and MAOM, and that these effects would be observed down to a 30 cm depth where root biomass remains sufficiently abundant.
2. MAOM would predominantly derive from microbial by-products, while POM would be more directly linked to plant inputs.
3. Tipping points in soil C pool variation along the disturbance gradient would occur at the transition from annual to perennial systems, as well as from herbaceous to woody vegetation.

## 2 Materials and Methods

### 2.1 Site description

The study was conducted in the floodplain of Lake Saint-Pierre (LSP), Québec, Canada (46,12°N, 72,49°W, alt. 6.2 m a.s.l). This region is categorized by a humid continental climate with an average daily temperature of 5.6 °C and an average annual precipitation of 948 mm (716 mm rain and 213 cm snow) based on 39-year data (1981-2020, Nicolet station) (Environment Canada, 2025). The LSP covers an area of around 50 000 ha and is recognized as a RAMSAR site (1998) and a UNESCO Biosphere Reserve (2000). The LSP is a widening of the Saint Lawrence River, Québec, Canada, formed by an immense rocky cavity partially filled by Quaternary marine clay sediments from the Champlain Sea, which gradually retreated around 12,000 years ago. The soil type of the floodplain is essentially composed of Gleysols, typical of flood-affected areas, with a poorly developed profile on sandy or clayey sedimentary material, covered by an alluvial loam with a high organic matter. This floodplain is driven by an annual spring flooding event following snowmelt, which lasts for 5-9 weeks and inundate croplands and surrounding natural ecosystems. Therefore, we refer to the floodplain as the relatively flat topographic area adjacent to Lake Saint-Pierre that experience flooding at biennial intervals (Watson et al., 2024). The total area of the flood recurrence zone is approximately 28,000 ha, indicating that the LSP floodplain is the largest wetland in the 1197-km-long Saint Lawrence River system (Hudon et al., 2018; Watson et al., 2024).

### 2.2 Experimental design

We established four study blocks corresponding to four areas of the littoral zone (floodplain) to represent regional differences (municipalities of Baie-du-Febvre [BAIE], Saint-Barthélemy [BART], La-Visitation-de-l’Île Dupas [DUPA] and Pierreville [PIER]) (Table 1). In each block, we were able to identify different land uses (all land uses were not necessarily present within the landscape of each block), ranging from the most to the least disturbed: 1) conventional croplands (maize [*Zea mays* L.]–soybean [*Glycine max* (L.) Merr.] alternation), 2) improved croplands (maize–soybean alternation with three-year-old intercropping and canary grass riparian strip), 3) temporary meadows (cut-grazed, established less than 5 years ago), 4) permanent meadows (cut-grazed, established more than 5 years ago), 5) marshes and 6) forested swamps (dominated by silver maple [*Acer saccharinum* L.] and black ash [*Fraxinus nigra* M.]). We studied a total of 30 sites, distributed as follows: eight conventional croplands (two in each block), eight improved croplands (two in each block), three temporary meadows (one in BAIE and two in BART), four permanent meadows (one in BAIE and BART and two in DUPA), three marshes (one in BAIE, BART and PIER), and four forested swamps (one in each bock) (see Campeau et al. (2024) for details).

**Table 1.** Means (±SEM) of soil characteristics at the 0-10 cm depth across all sites for the four studied blocks.

| Soil Variable | BAIE | BART | DUPA | PIER |
| --- | --- | --- | --- | --- |
| OM (%) | 10.44 ± 2.20 | 8.85 ± 0.86 | 6.67 ± 1.21 | 5.52 ± 1.68 |
| pH | 6.99 ± 0.18 | 5.70 ± 0.07 | 5.79 ± 0.15 | 5.62 ± 0.17 |
| Eh (mV) | 215.56 ± 6.34 | 270.25 ± 12.89 | 279.14 ± 11.21 | 289.42 ± 8.37 |
| Clay (%) | 32.61 ± 1.10 | 27.06 ± 1.75 | 20.67 ± 1.02 | 8.74 ± 1.23 |
| Silt (%) | 64.59 ± 1.23 | 62.62 ± 1.70 | 55.13 ± 2.77 | 49.76 ± 8.19 |
| Fe oxides (mg g<sup>-1</sup>) | 1.51 ± 0.17 | 1.29 ± 0.09 | 1.74 ± 0.13 | 0.66 ± 0.13 |
| Al oxides (mg g<sup>-1</sup>) | 1.07 ± 0.13 | 0.65 ± 0.08 | 0.55 ± 0.07 | 0.15 ± 0.06 |

OM: organic matter; pH: potential of hydrogen in soil measured in water; Eh: soil redox potential; Fe oxides: crystalline iron oxides; Al oxides: crystalline aluminum oxides. The blocks were: BAIE: Baie-du-Febvre; BART: Saint-Barthélemy; DUPA: La-Visitation-de-l'Île Dupas and PIER: Pierreville.

### 2.3 Measurement of vegetation biomass and Normalized Difference Vegetation Index (NDVI)

We measured aboveground and belowground vegetation biomass from August 16 to 29, 2021, during the peak of the vegetation period. Only roots with a diameter of 3 cm or less were sampled to ease extraction and minimize disturbance to vegetation growth in all land uses. For croplands, we measured the interrow spacing and sampled aboveground biomass along 1 m and 2 m rows for soybean and maize, respectively, while we sampled belowground biomass in pits matching the interrow width, with dimensions of 30 cm in length and 30 cm in depth. For meadows and marshes, we set up protected zones (30 × 30 cm) at the start of the growing season and harvested aboveground and belowground biomass to a depth of 30 cm. For forested swamps, we harvested litter and belowground biomass of trees from two pits per site, each measuring 25 × 25 cm in surface area and 30 cm in depth. All fresh biomass collected from each land use was oven-dried at 80°C for 3 days and then weighed to calculate the dry biomass by the sampled area. In addition, we measured aboveground biomass of forests as the mean aboveground biomass of trees multiplied by their mean density. We determined the mean forest density using the mean point-tree distance method at a series of 10 stations spread over a 500 m distance (Elzinga et al 1998). We calculated the mean aboveground biomass (kg) by measuring 20 trees and using the allometric equation Eq. (1):

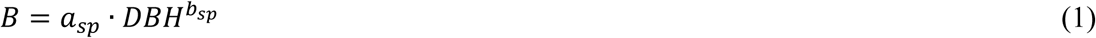

Where *B* is aboveground biomass, *DBH* is diameter at breast height and *a_sp_* and *b_sp_* represent the parameters associated with the two dominant tree species, *A. saccharinum* (*a* = −2.047; *b* = 2.3852) and *F. nigra* (*a* = −2.0314; *b* = 2.3524) (Chojnacky et al., 2014).

We also measured NDVI, which is another metric of terrestrial vegetation density (Huang et al., 2021). NDVI values were obtained from the European Union’s Copernicus Land Monitoring Service information (2024), providing 10-day NDVI composites at 300 m resolution for our study area. For each site, NDVI was averaged over the period from May to September 2021.

### 2.4 Soil sampling and analyses

All 30 sites were sampled at the same elevation of 6.2 m a.s.l to ensure the same flooding frequency. At each site, we randomly collected three soil cores, each measuring 1m in depth and 5.1 cm in diameter, using a thermal drill and plastic tubes in October 2019 (AMS Gas Powered REDI Boss Hammer, American Falls). We then sliced the cores into 12 sections at the following depth intervals: 0-5, 5-10, 10-15, 15-20, 20-25, 25-30, 30-35, 35-40, 40-50, 50-60, 60-80, and 80-100 cm. The corresponding slices from the three cores were pooled to obtain one composite sample for each depth interval at each site and air-dried. After acid fumigation, we measured organic C and total nitrogen (N) concentrations of all composite samples, along with their δ¹³C values using an elemental analyzer coupled with an isotope ratio mass spectrometer (EA-IRMS, technology Agilent, Santa Clara, CA, USA). We sieved 10 g subsamples to 1 mm and oven-dried them at 80°C for 36 hours to estimate the dry mass of samples. We calculated soil *Bulk density* (g cm^-3^) using the equation Eq. (2) (Throop et al., 2012):

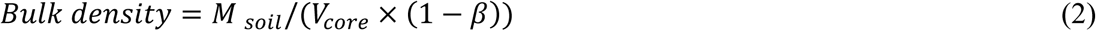

Where *M_soil_* is the mass (g) of an oven-dried sample, *V_core_* is the volume of the soil core (cm^3^) and *β* is the volume fraction of pebbles and coarse plant debris (%/100), using densities of 2.65 and 0.7 g cm^-3^, respectively (Don et al., 2007). Finally, we calculated *SOC stock* (Mg ha^-1^) as follows (Eq. (3)):

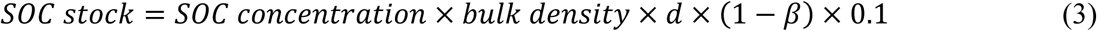

Where *SOC concentration* is the soil organic carbon concentration (g kg^-1^), *d* is the depth (cm) and *β* is the volume fraction (%/100) of pebbles and coarse plant debris. We calculated cumulative SOC stocks at a given depth by summing the SOC stocks at that depth and those above it.

Furthermore, we also carried out soil samplings (0-10 cm depth) at the 30 sites on five occasions (October 2019, June and October 2020, October 2021, and November 2022) for microbial biomass and dissolved organic carbon (DOC) analyses. We brought these soil samples to the laboratory in a cooler, sieved them to 2 mm and stored them at −80°C. We then conducted a chloroform fumigation-extraction using 0.5M K_2_SO_4_ on these samples to determine microbial biomass (Vance et al., 1987). Non-fumigated samples were used to quantify DOC, while microbial biomass was calculated from the difference between fumigated and non-fumigated samples. The DOC analyses were performed using a portable M9 TOC analyzer (Sievers, SUEZ).

### 2.5 Soil fractionation

Particle-size fractionation by wet sieving was carried out on 10 g of air-dried soil from each sample to study SOC stabilization. Each 10 g subsample was placed into a 250 ml plastic bottle and agitated in a solution of sodium hexametaphosphate (0.5 %) with 10 glass beads (6 mm diameter) overnight under continuous and regular movement to completely disperse soil particles (Balesdent et al., 1991). The soil fractions that passed through the 53 μm sieve and those remaining on the sieve were defined as the mineral-associated (< 53 μm) and the particulate (≥ 53 μm) fractions, respectively. We determined the concentration of mineral-associated organic matter - C (*MAOM-C_concentration,_* g kg^-1^ fraction) and particulate organic matter - C (*POM-C_concentration_*), using the same method as for SOC concentration in the bulk soil. To account for the distribution of each fraction in the bulk soil, we used the amount of *MAOM-C* or *POM-C* (g kg^-1^ soil) in this study, which was calculated using the equation Eq. (4) (d’Annunzio et al., 2008):

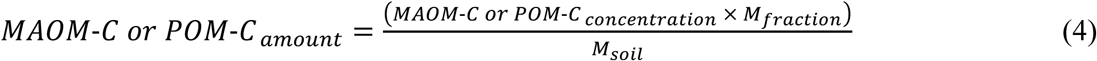

where *MAOM*-*C or POM*-*C_concentration_* is the concentration of mineral-associated organic C or particulate organic C (g kg^-1^ fraction), *M_fraction_* is the dry mass of the soil mineral-associated or particulate fraction, and *M_soil_* is the dry mass of the bulk soil used for fractionation, using soil moisture at 80 °C.

### 2.6 Determination of plant-derived C

Using the fact that δ^13^C in the subsoil for all land uses converged towards the same value (see Results section), which can be considered as the reference or old δ^13^C of past vegetation, and that δ^13^C in the topsoil tended towards that of current vegetation (Table S1), we were able to estimate the C derived from current vegetation in the topsoil (Balesdent and Mariotti, 1996). It should be noted that we had to weight-average the δ^13^C of maize and soybean to obtain the plant δ^13^C in both croplands (Table S1). For each land use, we calculated the C derived from current vegetation in each soil fraction (*Plant-derived C_MAOM or POM_*, g kg^-1^ soil) using the equation Eq. (5) (Balesdent and Mariotti, 1996; Tonucci et al., 2017):

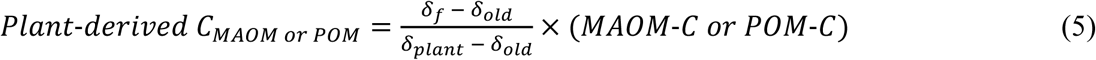

Where *MAOM-C* or *POM-C* was the amount of C in MAOM or POM, *δ_f_* was the δ^13^C of MAOM or POM, *δ_old_* was the reference or old δ^13^C values in the subsoil and *δ_plant_* was the δ^13^C of current vegetation.

### 2.7 Neutral and amino sugar analyses

We determined neutral and amino sugars from air-dried and ground (< 0.15 mm) soil subsamples and fine fractions (MAOM) using the method detailed in Chantigny and Angers (2008). Briefly, we extracted neutral sugars using H₂SO₄ solution, neutralized the hydrolysates with NaOH solution, sequentially passed the extractant through anion- and cation-exchange resins, and analyzed the purified solutions for plant-(arabinose and xylose; AX) and microbe-derived (galactose and mannose; GM) sugar concentrations (Gunina and Kuzyakov, 2015) using high-performance anion-exchange chromatography (Model Dionex ICS 5000+, Thermo Scientific, Sunnyvale, CA).

To estimate fungal and bacterial amino sugars, we quantified glucosamine (GluN) and muramic acid (MurA), respectively. Briefly, we extracted amino-sugars using HCl solution and N_2_ bubbling and analyzed their concentration by high-performance liquid chromatography (Model Dionex ICS 5000+, Thermo Scientific, Sunnyvale, CA). The amounts of each sugar in MAOM were calculated based on the C mass present in that sugar and by taking into account the distribution of the fine fraction in the bulk soil (i.e., mg C g^−1^ soil). Sugar amounts in POC were estimated by subtracting sugar amounts in MAOC from that in the bulk soil.

### 2.8 Statistical analysis

All statistical tests were performed using R software version 4.4.0 and RStudio version 2024.04.0 (R Development Core Team, 2013). We produced all figures using the visreg and ggplot2 packages of R software (Breheny and Burchett, 2017).

#### 2.8.1 Comparison of biomass and the different forms of SOC between land uses

We first compared plant aboveground and belowground biomass, NDVI, and soil microbial biomass among land uses using linear mixed-effect models (LMMs), where ‘site’ was a random effect (lme4 and lmerTest R packages; Bates et al., 2015; Kuznetsova et al., 2017). Model assumptions (independence of residuals, homoscedasticity and normality of residuals) were assessed using diagnostic graphs and Shapiro-Wilk tests. We applied log transformations to aboveground, belowground and soil microbial biomass to meet the model assumptions. We used Tukey’s Honest Significant Difference (HSD) test as a *post-hoc* method to make pairwise comparisons where appropriate using “emmeans” function (emmeans R package; Lenth et al., 2018).

We then assessed the relationship between soil depth and SOC concentration, SOC stock, BD or soil δ^13^C across land uses to obtain an overview of these variables along the soil profiles (0-100 cm). We used generalized linear mixed-effects models (GLMMs) to produce trend curves for SOC concentration, SOC stock and BD using gamma distributions (log, log, and identity link, respectively) in the “glmer” function (lme4 R package; Bates et al., 2015), and including land use and log-transformed depth as fixed effects since model assumptions were not met (Hengl and MacMillan, 2019). We fitted LMMs for δ^13^C after applying a square-root transformation of δ^13^C + 30 and a log transformation of soil depth. For all these models, site and sample core were included as random effects to account, respectively, for site effects and the dependence of SOC values across soil depths. We also analyzed these soil variables at each soil depth separately to determine the depth of land use effects (Table S2); for these models, LMMs with ‘site’ as a random effect met assumptions without transformation.

Based on the results of the comparison above, we conducted a more detailed analysis (LMMs and pairwise comparisons) only at the 0-10 cm for SOC concentration, SOC stock, DOC, POM-C, and MAOM-C, following the same method as for plant biomass. We included clay + silt percentage as a predictor in the SOC stock model and clay percentage in the MAOM model as fine particles are considered to be an important factor for SOC retention on soil minerals and within soil aggregates (Six et al., 2002; Prout et al., 2021). We applied log transformations for POM-C to meet the model assumption.

#### 2.8.2 Assessment of the origins of stable SOC

We compared δ¹³C, plant-derived C, plant-, microbial-, fungi-, and bacteria-derived sugars in MAOM and POM at the 0–10 cm soil depth between land uses following the same method as for plant biomass. We applied log transformations for plant-derived C and plant-, microbe-, and bacteria-derived sugars in MAOM and POM to meet model assumptions. To test our hypothesis, we also compared microbe-versus plant-derived sugars, and fungi-versus bacteria-derived sugars, in MAOM and POM using LMMs.

#### 2.8.3 Assessment of variation in all soil C pools along a disturbance gradient

We conducted a principal component analysis (PCA) to obtain an overview of all soil C pools at the 0–10 cm depth across land uses using the FactoMineR R package. Variables included SOC concentration, MAOM-C, POM-C, plant-derived C, plant- and microbe-derived sugars in bulk soil, and soil microbial biomass. We then correlated the first PCA axis, which explained 83% of the overall variation, with the disturbance gradient using a local polynomial regression (LOESS) via “geom_smooth” function of the ggplot2 R package to capture non-linear relationships. This gradient was created by assigning numeric values at a constant interval, using the “as.numeric” base R function, to each land use arranged from the most to the least disturbed: (1) conventional croplands, (2) improved croplands, (3) temporary meadows, (4) permanent meadows, (5) marshes, and (6) forested swamps. For clarity, numeric values were subsequently replaced with the corresponding land use names.

## 3 Results

### 3.1 Vegetation and soil microbial biomass and NDVI

Forested swamps had much higher aboveground biomass compared to the two croplands, the two meadows (temporary and permanent) and marshes (*P* < 0.001, Fig. 1a). Forested swamps, marshes and permanent meadows had higher belowground (*P* < 0.001) and soil microbial (*P* < 0.001) biomass than the two croplands and temporary meadows (Figs. 1cd). NDVI values were higher in forested swamps than in conventional croplands (*P* < 0.01, Fig. 1b).

**Fig. 1.**
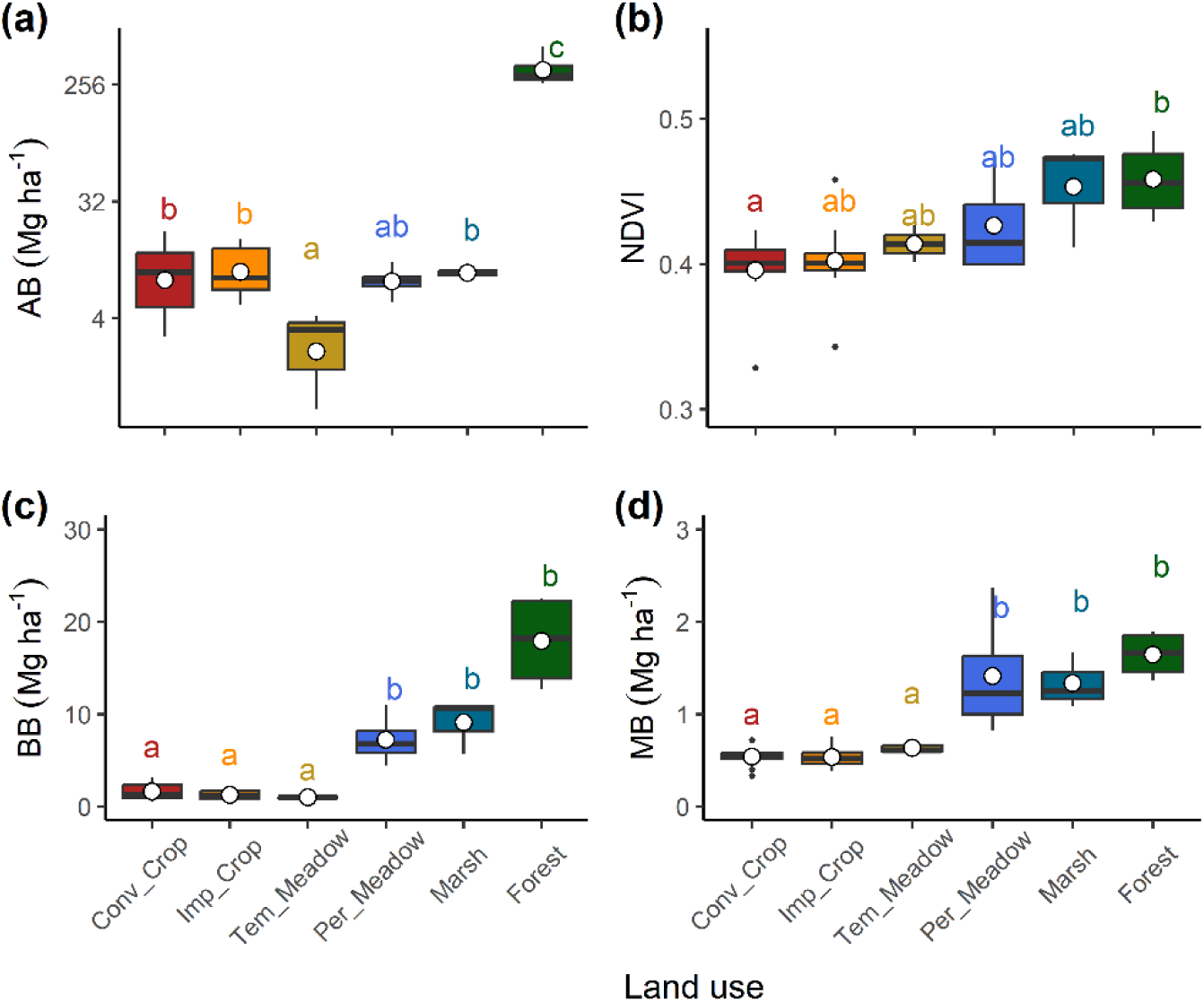
Aboveground (AB) (a), belowground (BB) (c) and soil microbial (MB) (d) biomass, and Normalized Difference Vegetation Index (NDVI) (b) among studied land uses. Belowground biomass was collected at the 0-30 cm depth, while soil microbial biomass was estimated at the 0-10 cm depth. NDVI was calculated as the mean of NDVI values from May to September. Different lowercase letters indicate significant differences between land uses (*P* < 0.05). Conv: conventional; Imp: improved; Per: permanent; Tem: temporary.

### 3.2 Soil organic carbon, bulk density and δ13C along the soil profile

The interaction between land use and soil depth significantly affected SOC concentration and stock, BD, and δ^13^C along the soil profile of 0-100 cm (*P* < 0.001 for all, Fig. 2). For each depth separately, SOC concentration was influenced by land use down to the 10 cm depth and consistently decreased with increasing soil depth (Fig. 2a). Land use impacted SOC stock only at the 0-5 cm depth (*P* < 0.01, Fig. 2c), whereas it influenced cumulative SOC stock at the 0-10 cm depth (P=0.02, Table S2). In addition, land use had a significant impact on BD down to the 20 cm depth (Fig. 2b). The effect of land use on soil δ^13^C was highly significant down to the 25 cm depth, but its effects were also observed at the 50-80 cm depth (Fig. 2d). Across all land uses, the soil δ^13^C values converged towards −25.5‰ with increasing depth, aligning with the mean soil δ^13^C at the 40-100 cm depth (Fig. 2d).

**Fig. 2.**
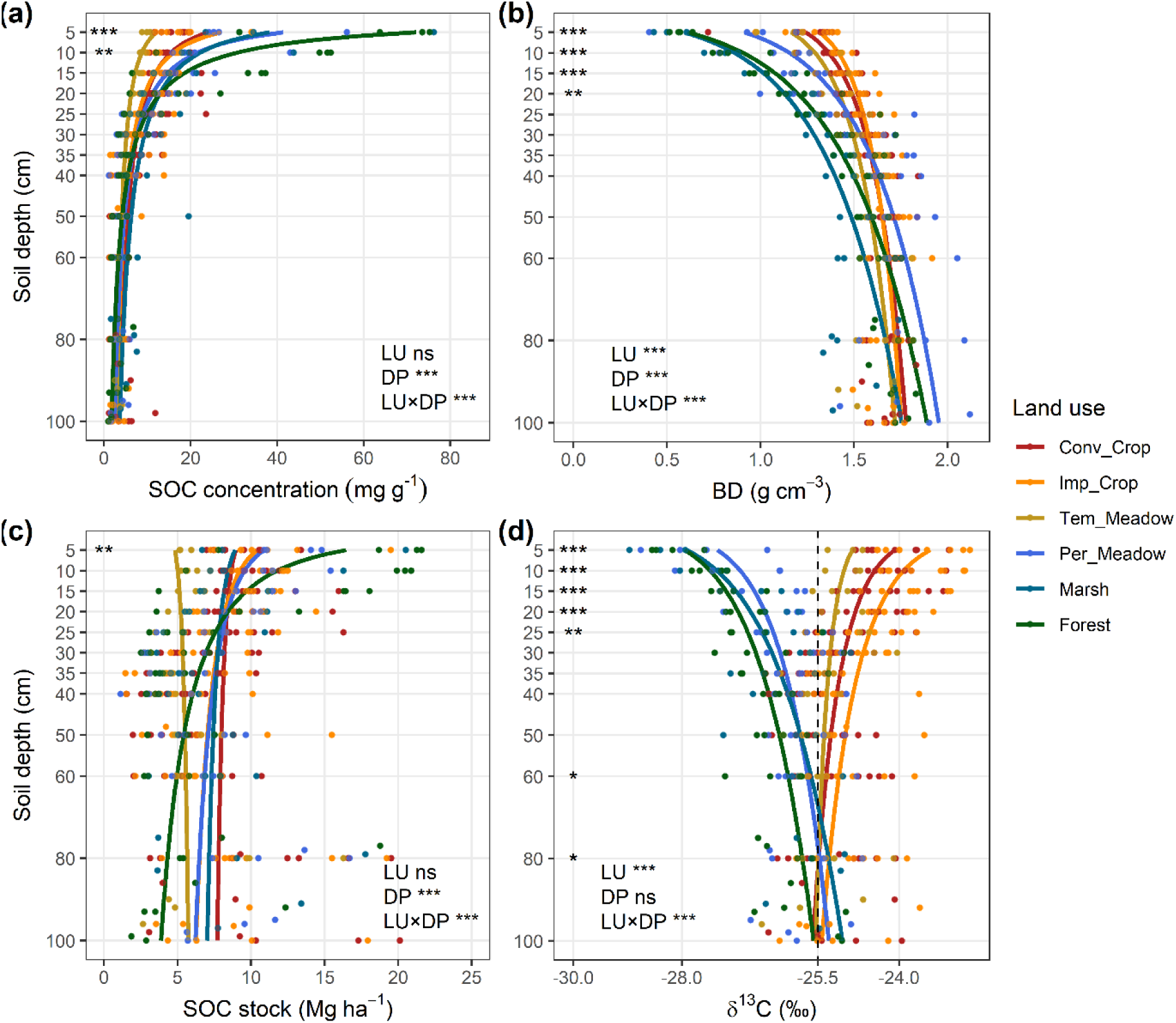
Soil organic carbon (SOC) concentration (a), bulk density (BD) (b), SOC stock (c), and soil δ^13^C (d) as a function of depth for the different land uses. Trend curves were obtained from mixed models with land use (LU) and soil depth (DP) as predictors. Significance code: ***: *P* < 0.001; **: *P* < 0.01; *: *P* < 0.05; ns: nonsignificant. Stars to the right of a depth indicate significant differences between land uses at that depth, considered separately (See Table S2 for complete details). The dotted line in panel d represents the mean δ^13^C in the deep soil layer of 40-100 cm (−25.5‰), a value considered to be the reference or old δ^13^C. Conv: conventional; Imp: improved; Per: permanent; Tem: temporary.

### 3.3 Comparison between land uses of different forms of organic C at the 0-10 cm soil depth

Forested swamps had higher SOC concentrations than both types of croplands and temporary meadows at the 0-10 cm depth (Fig. 3a) and had higher cumulative SOC stock than temporary meadows at the 0-10 cm depth (Fig. S1). Clay + silt percentage positively affected SOC stock (*P* < 0.01, estimate = 0.18) at this depth. Mineral associated-organic C (MAOC) was affected by land use (*P* < 0.01) and was positively correlated with clay percentage (*P* = 0.01, estimate = 0.50) at this depth. The amount of MAOM-C was greater in forested swamps than in both types of croplands (Fig. 3d). The MAOM-C amounts for all floodplain land uses remained below the maximum observed capacity (saturation) of 87 mg C g^−1^ clay + silt (Fig. 3e; Georgiou et al., 2025). Overall, MAOM-C varied less among land uses than POM-C (Figs. 3cd). Land use also affected POM-C at the 0-10 cm depth (*P* < 0.001), with higher amount in forest swamps, marshes and permanent meadows than in temporary meadows and both croplands (Fig. 3c). In addition, land use had significant effects on DOC (*P* < 0.001), with forested swamps showing higher DOC than the two croplands and the two meadows at this depth (Fig. 3b).

**Fig. 3.**
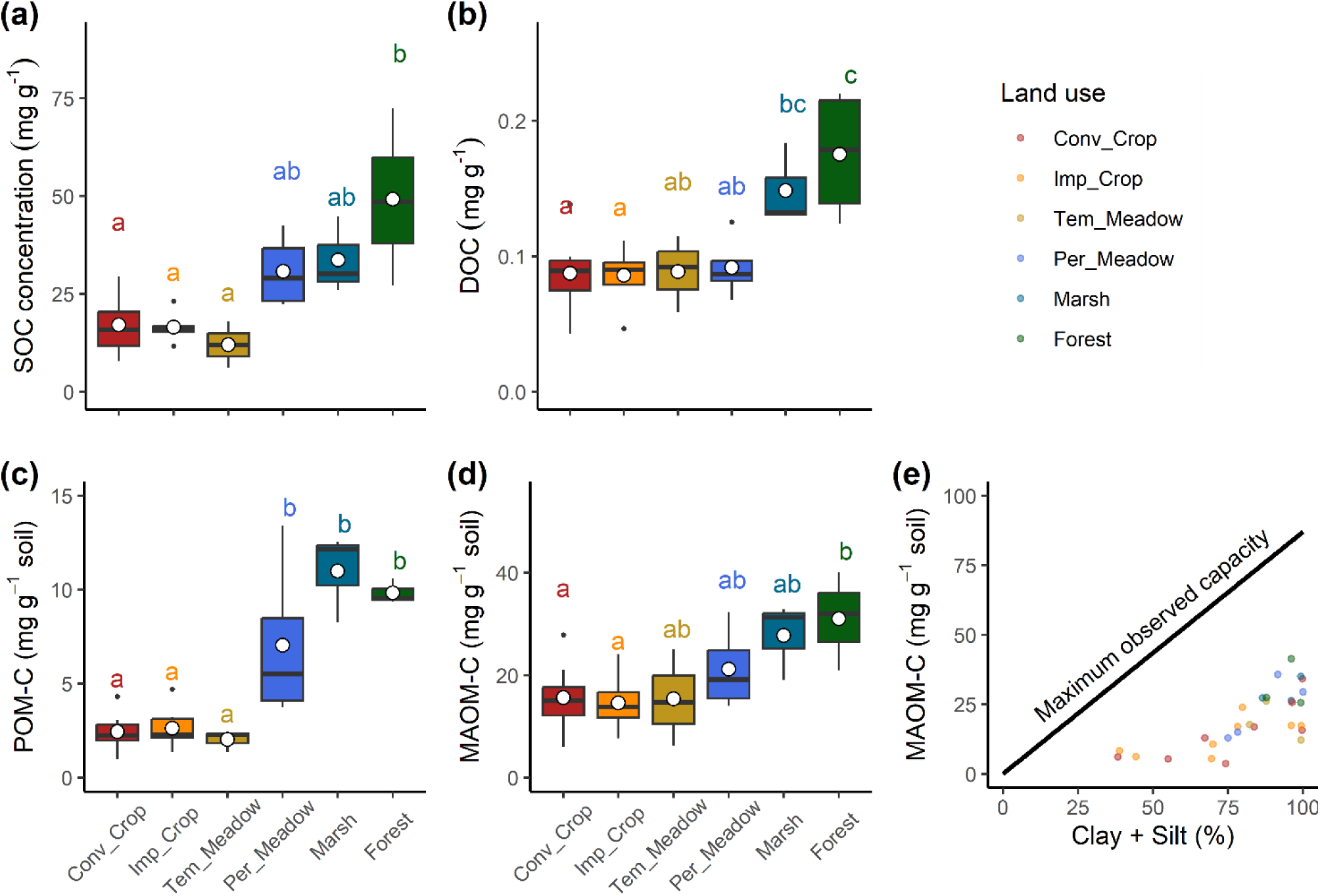
The effects of land use on the different forms of organic C at the 0-10 cm soil depth and the maximum observed capacity for MAOM-C storage. (a) Soil organic carbon (SOC) concentration; (b) dissolved organic carbon (DOC), (c) particulate organic carbon amount (POM-C), (d) mineral-associated organic carbon amount (MAOM-C), (e) MAOM-C as a function of clay + silt percentage at the 0-10 cm depth, with the maximum observed capacity (87 mg C g^−1^ clay + silt) for MAOM-C storage according to Georgiou et al. (2025). Different lowercase letters indicate significant differences between land uses (*P* < 0.05). Conv: conventional; Imp: improved; Per: permanent; Tem: temporary.

### 3.4 Origin of stable SOC at the 0-10 cm depth

Land use significantly impacted δ^13^C in both MAOM and POM (P < 0.001 for both). Compared to the δ^13^C values of POM, those of MAOM deviated less from the reference or old δ^13^C (–25.5‰), which represents the mean δ^13^C in the deep soil layer (Figs. 4ab). Only the δ^13^C values of MAOM in forested swamps (–28.23‰), marshes (–28.16‰), and permanent meadows (–27.51‰), deviated from this reference, but not those in croplands and temporary meadows (Fig. 4a). The δ^13^C values of POM deviated from this reference toward lower values in forested swamps (– 29.33‰), marshes (–28.79‰), and permanent meadows (–28.35‰), whereas those from the conventional (–22.66‰) and improved (–21.64‰) croplands shifted toward higher values (Fig. 4b). Plant-derived C in MAOM and POM was significantly different between land uses at the 0-10 cm depth (*P* < 0.01 for both). More plant-derived C was present in MAOM in the forested swamps and marshes than in both types of croplands, and there was more plant-derived C in POM in the forested swamps than in both types of croplands and temporary meadows (Fig. 4cd).

**Fig. 4.**
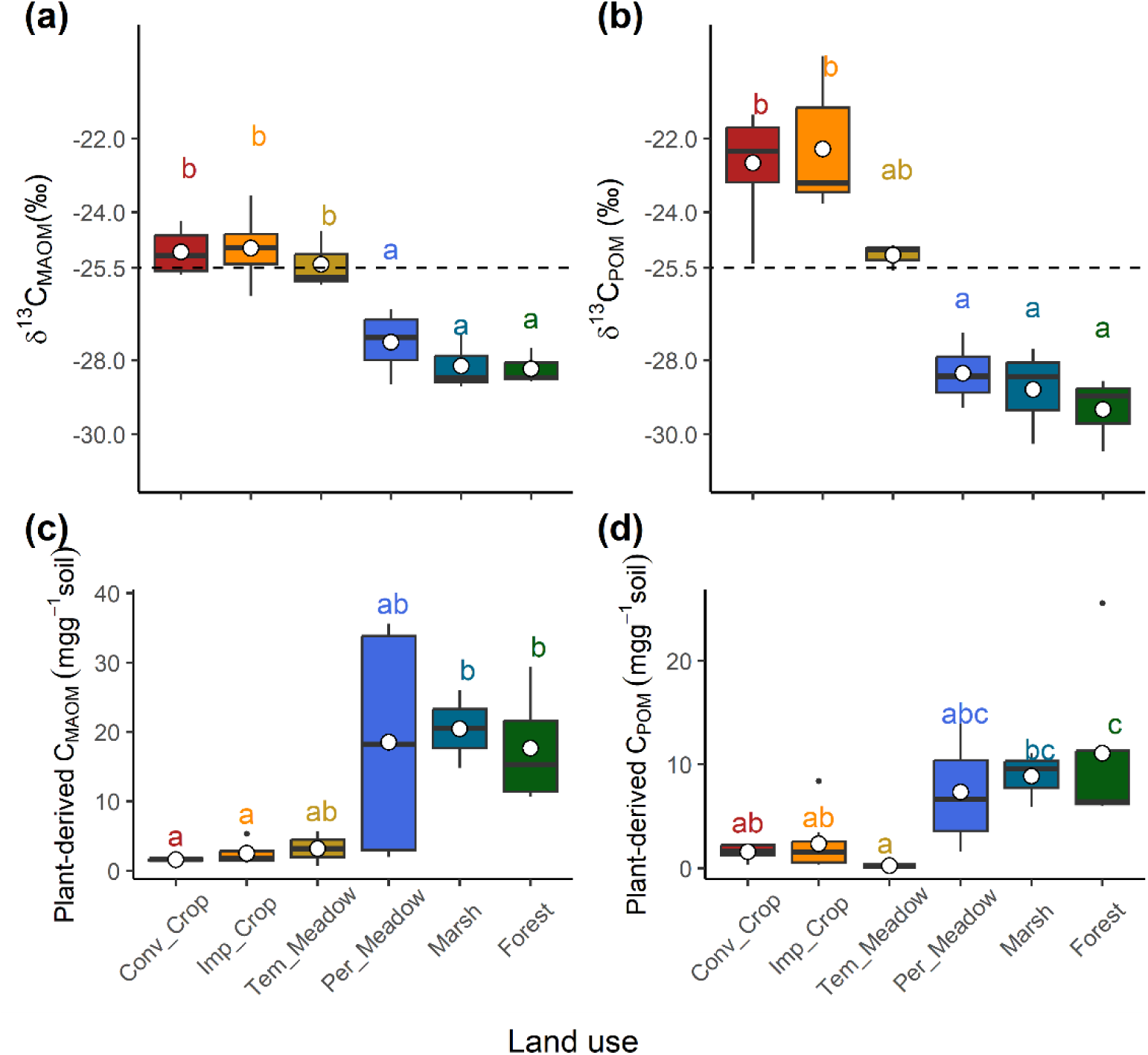
δ^13^C values and plant-derived C in mineral-associated organic matter (MAOM) (a,c) and in particulate organic matter (POM) (b,d) across land uses at the 0-10 cm depth. Plant-derived C was estimated using the δ^13^C tracing technique (see the section 2.6 for details). The dotted line represents the reference δ^13^C (−25.5‰). Different lowercase letters indicate significant differences between land uses (*P* < 0.05). Conv: conventional; Imp: improved; Per: permanent; Tem: temporary.

Microbe-derived sugars were more abundant in MAOM than plant-derived sugars (P < 0.01), whereas the opposite trend was observed in POM (P < 0.05; Fig. 5). Forested swamps and marshes had higher amounts of microbe- and plant-derived sugars in MAOM and POM compared to the two croplands (P < 0.001 for all; Fig. 5). Within soil microbes, fungi-derived sugars were more abundant than bacteria-derived sugars in both MAOM and POM (P < 0.001 for all; Fig. 5). Forested swamps had higher fungi-derived (P < 0.01) and bacteria-derived (P < 0.001) sugars in MAOM than both types of croplands, while permanent meadows exhibited higher fungi-derived sugars in POM (P < 0.001) than both types of croplands (Fig. 5).

**Fig. 5.**
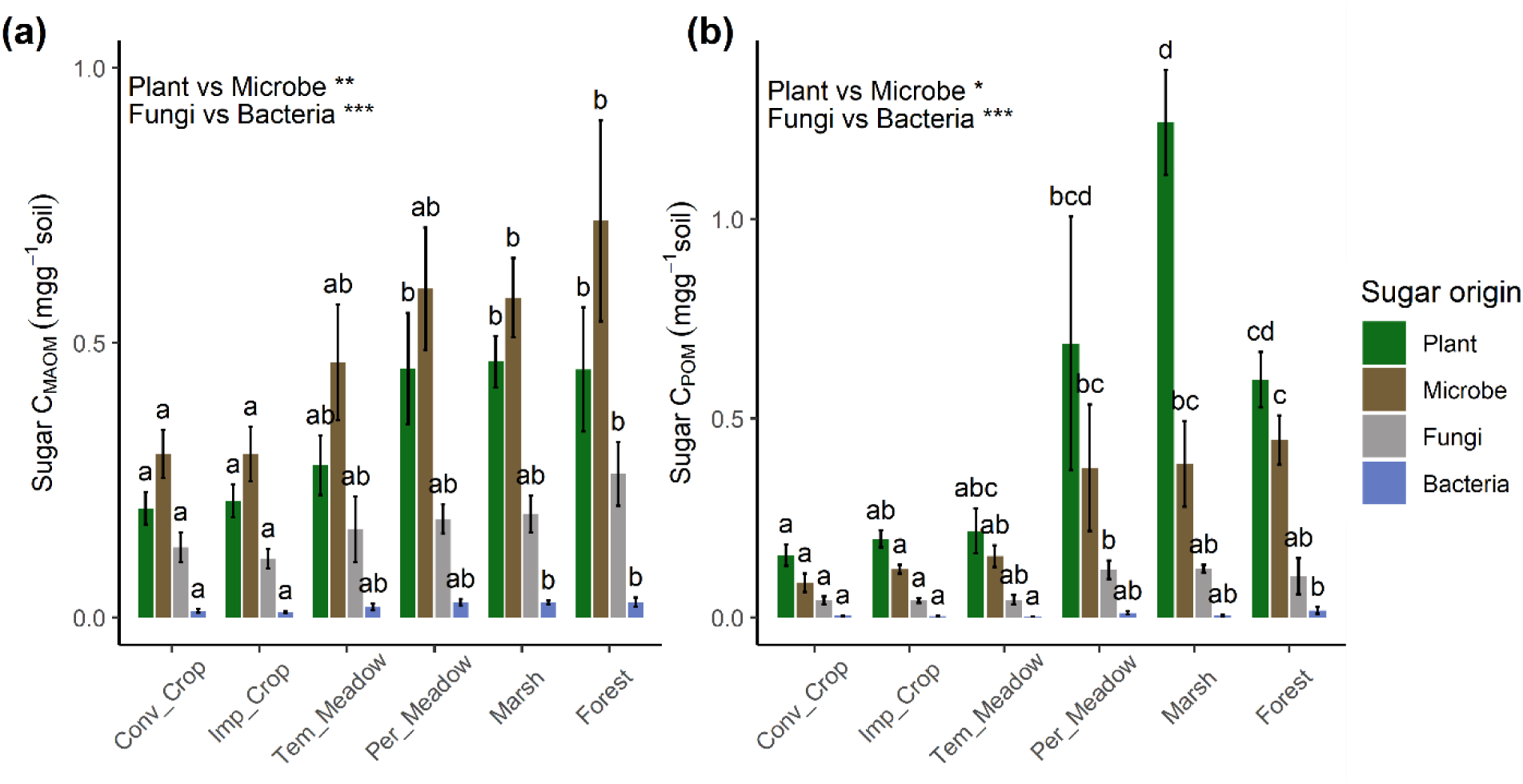
Sugar origin in mineral-associated organic matter (MAOM) (a) and in particulate organic matter (POM) (b) across land uses at the 0-10 cm depth. Different lowercase letters indicate significant differences between land uses (*P* < 0.05). Stars indicate significant differences between plant- and microbe-derived sugars or between fungi- and bacteria-derived sugars: ***: *P* < 0.001; **: *P* < 0.01; *: *P* < 0.05. Sugar amounts were calculated based on the C mass present in each sugar and by taking into account the distribution of each soil fraction in the bulk soil (i.e., mg C g^−1^ soil). Conv: conventional; Imp: improved; Per: permanent; Tem: temporary.

### 3.5 Variation in soil C pools along a disturbance gradient

A clear distinction of all soil C pools was observed between temporary (croplands and temporary meadows) and permanent land uses (permanent meadows, marshes, and forested swamps), as shown by PCA (Fig. 6a). The overall soil C pools, represented by PCA1, increased with decreasing anthropogenic disturbance intensity, ranging from conventional croplands to forested swamps (Fig. 6b). However, these soil C pools did not follow a linear pattern along the disturbance gradient, with a tipping point occurring at the transition between temporary and permanent meadows (Fig. 6b).

**Fig. 6.**
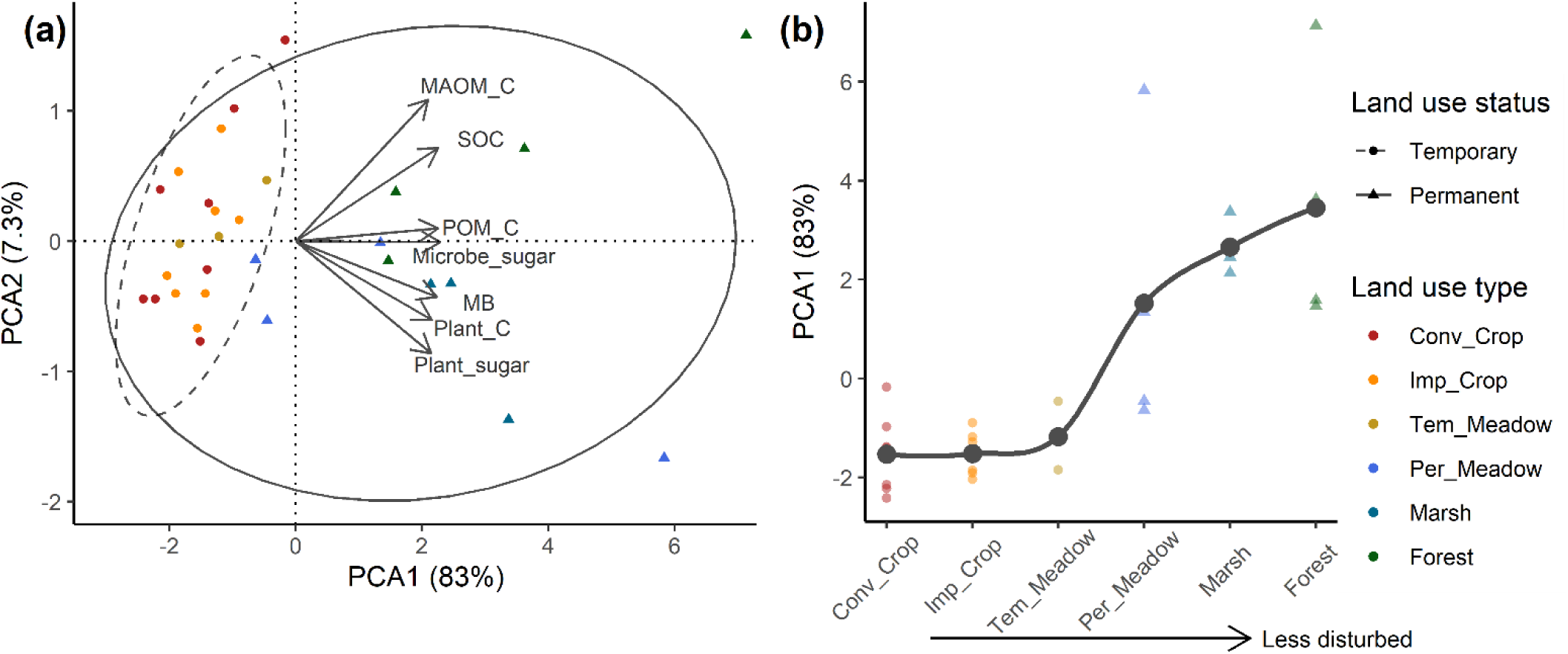
Principal component analysis (PCA) of all soil carbon (C) pools (a) and variation of overall soil C pools along the anthropogenic disturbance gradient at the 0-10 cm depth (b). Conv: conventional; Imp: improved; MAOM-C: mineral-associated organic C; MB: soil microbial biomass; Per: permanent; POM-C: particulate organic C; SOC: Soil organic C: Tem: temporary.

## 4 Discussion

### 4.1 Key role of land uses in SOC storage in floodplain topsoil

The effects of land use on SOC concentration and stock in the LSP floodplain were mostly observed in the top 10 cm, partially aligned with our hypothesis. Forested swamps had higher SOC and DOC than croplands in our floodplain topsoil, consistent with previous studies (Omengo et al., 2016; D’Elia et al., 2017), likely due to high above- and belowground biomass inputs. Notably, root C may be retained more efficiently in SOC than aboveground inputs, as the disproportionate difference in aboveground biomass between forested swamps and marshes was not reflected in SOC stocks (Ghafoor et al., 2017; Sokol et al., 2019; Rabearison et al., 2025). In addition, previous studies found that autochthonous OC from local vegetation is the main source of SOC in floodplain topsoil, outweighing sediment inputs, as demonstrated by δ¹³C and ¹⁴C analyses (Omengo et al., 2016; Scott and Wohl, 2018). For both croplands, low biomass inputs combined with repeated soil tillage likely explain the low SOC storage in the topsoil (Bronick and Lal, 2005; Ciais et al., 2010).

Land use effects on SOC stock were no longer evident below 10 cm depth, since vegetation inputs were found to have small contributions to SOC in floodplain subsoils (Omengo et al., 2016). The low bulk density in forested swamps and marshes, compared to croplands, likely also reduced differences in subsoil SOC stock. However, the soil δ¹³C signature indicates that land use and the associated vegetation inputs influenced SOC composition down to 25 cm but had little impact below 40 cm depth. Frequent flood events could cause partial loss of these vegetation inputs, limiting OM accumulation in deeper soil layers (Saint-Laurent et al., 2017).

### 4.2 Is stable SOC dominated by microbial- or plant-derived C?

Soil in forested swamps contained more C as MAOM than cropland soils, highlighting the role of trees in promoting accumulation of stable SOC in the LSP floodplain (Del Galdo et al., 2003; Shoumik et al., 2025). In line with our hypothesis, microbe-derived sugars were more abundant in MAOM than plant-derived sugars, indicating that microbial inputs are the primary C source in MAOM. The high biomass inputs from forested swamps would thus have been efficiently utilized by soil microbes, enhancing microbial by-products and MAOM-C accumulation (Cotrufo et al., 2013; Liang et al., 2017). As in van Leeuwen et al. (2017), higher microbial biomass in forest topsoil, relative to croplands, was found in our study, supporting the involvement of microbial metabolism in the SOC stabilization mechanism. Microbial by-products, particularly necromass, often have a stronger affinity for mineral phases than plant-derived compounds and may be less accessible to decomposers (Kleber et al., 2021; Zhao et al., 2025). Within microbial communities, the higher fungi-derived sugars found in MAOM under all land uses, compared to bacteria-derived sugars, highlights the important contribution of fungi in the formation of stable SOC (Fu et al., 2025).

The δ¹³C and sugar analyses also confirm that some plant-derived inputs accumulated directly in MAOM without microbial assimilation. In forested swamps, marshes and permanent meadows, the MAOM δ¹³C values shifted towards those typical of trees and herbaceous plants, indicating substantial inputs of plant-derived C in MAOM. These inputs are likely mediated by roots, as supported by the high belowground biomass under these land uses. Previous studies reported that soluble compounds derived from roots (exudates and rhizodeposits) can be directly adsorbed onto soil mineral surfaces, contributing significantly to MAOM (Hagedorn et al., 2015; Angst et al., 2017; Córdova et al., 2018). Plant inputs may also first undergo *ex vivo* transformation by microbial extracellular enzymes, producing labile plant-derived compounds that readily interact with soil minerals (Liang et al., 2017; Sokol et al., 2019; Whalen et al., 2022). Thus, enhancing plant-derived C in MAOM could be achieved by promoting land uses that favor labile compounds, such as herbaceous-rich land uses.

High clay and silt contents facilitated SOC storage and MAOM-C accumulation in our floodplain topsoil, as reported by several floodplain studies (Bullinger-Weber et al., 2014; Sutfin and Wohl, 2016; Heger et al., 2024). These fine-textured particles promote the adsorption of organic C onto mineral and metal oxide surfaces, leading to the formation of stable organo-mineral complexes or MAOM (Cotrufo et al., 2013; Lavallee et al., 2020). Regardless of land use, MAOM was not saturated in the LSP floodplain, compared to the maximum storage capacity of 87 mg C g^-1^ clay + silt proposed by Georgiou et al. (2025). Therefore, there could be considerable potential to stabilize further SOC in this area.

### 4.3 Why does POM remain important?

Our sugar and δ¹³C analyses indicated that plant-derived compounds were the dominant source of C in POM. Accordingly, many studies have shown that POM is mainly comprised of partially decomposed OM from vegetation (Balesdent, 1996; Six et al., 2002). Significantly more POM-C was found in forested swamps, marshes and permanent meadows than in croplands and temporary meadows, highlighting the importance of this SOC reservoir in soils under forest and permanent herbaceous-dominated vegetation (Yeasmin et al., 2020). Recent findings suggest that POM constitutes a key compartment for additional C storage as it may not be subjected to saturation mechanisms (Dignac et al., 2025; Georgiou et al., 2025). The greater contrast in POM-C than MAOM-C found between croplands and permanently covered soils indicates that POM remains more vulnerable to anthropogenic disturbance than MAOM (Cotrufo et al., 2019; Angst et al., 2021).

We also detected microbial-derived compounds in POM, suggesting that some microbial by-products remain unprotected by mineral surfaces. During decomposition, POM is not entirely lost as CO_2_, but a portion can subsequently be transferred into MAOM through microbial metabolism (Liang et al., 2017; Dignac et al., 2025). Thus, increased POM could serve as a potential reservoir for the eventual MAOM formation upon binding to mineral surfaces (Dignac et al., 2025; Georgiou et al., 2025). In this sense, it is important to promote land uses that enhance this C storage from POM to MAOM, such as forested swamps (Kallenbach et al., 2019).

### 4.4 Implications for land use management in floodplain ecosystems

The overall soil C pools were predominantly higher under land uses with permanent vegetation (permanent meadows, marshes, and forested swamps), emphasizing the critical role of conserving these stable ecosystems within the LSP floodplain. The variation of SOC pools was not linear along the disturbance gradient, with a tipping point occurring only in the transition from temporary to permanent meadows, but not during the shift from permanent herbaceous to woody vegetation (Figs. 6b and S2). In this context, prioritizing the conversion of temporary meadows or low-productivity croplands to permanent meadows, once prevalent in the early 1980s, is crucial (Yeasmin et al., 2020). Its establishment is a gradual process, likely requiring over five years to achieve significant SOC gains. For instance, the permanent meadow in Baie-du-Febvre has been maintained by the same family for at least three generations, according to the owner, and showed SOC comparable to those in forested swamps. Nevertheless, such conversion efforts must be accompanied by targeted outreach to address negative perceptions among floodplain farmers, particularly regarding the forage quality and competitive nature of the common weed bluejoint reedgrass (*Calamagrostis canadensis* [Michaux] Beauvois) (Sparrow and Panciera, 2005; Campeau et al., 2024).

As forested swamps were the most effective systems for the storage of stable SOC, we also recommend expanding their area through afforestation in low-productivity croplands while minimizing soil disturbance. Specifically, reforestation with silver maple in the LSP floodplain could not only increase SOC stabilization but also support maple syrup production, as this species is already dominant in the LSP landscape (Watson et al., 2024) and two silver maple sugarbushes equipped with tubing were observed in Pierreville. Fast-growing trees such as poplar (*Populus* spp.) may also be planted to promote rapid timber production and enhance SOC stabilization, as they tend to transfer more labile compounds that favor microbial by-product formation and hence MAOM accumulation (Cotrufo et al., 2013; Poirier et al., 2018; Rabearison et al., 2025). In addition, it should be noted that neither the agri-environmental practices (improved croplands) nor conventional croplands were beneficial for SOC stabilization in the LSP floodplain.

## 5 Conclusion

Land use significantly influenced SOC storage and stabilization in the topsoil of the LSP floodplain. Forested swamps had the highest vegetation and soil microbial biomass, as well as SOC and MAOM-C, highlighting their key role in long-term SOC stabilization. Permanent meadows and marshes also contributed significant C to POM, which appears to be an important SOC reservoir in herbaceous-dominated systems. Our study also showed that microbe-derived C was more abundant in MAOM, while plant-derived C dominated POM, providing a basis to improve SOC persistence predictions and guide targeted land use management. Decreasing anthropogenic disturbance enhanced SOC gains, with a tipping point occurring during the transition from temporary to permanent meadows. Therefore, conserving and expanding land uses with permanent vegetation covers in the LSP floodplain likely provides ecological benefits and supports climate change mitigation. Future studies on greenhouse gas (GHG) fluxes (CO_2_ and N_2_O) would further clarify net GHG sequestration in this landscape.

## Acknowledgements

We are grateful to the many people who allowed us access to their property, especially the farmers. This was arranged through an agreement with le pôle d’expertise multidisciplinaire en gestion durable du littoral du Lac Saint-Pierre. We thank Maxime Tremblay, Anne-Marie Decelles and Chantal Fournier, who provided administrative support. We thank Steven Tessier, Roxanne St-Pierre, Saylena Fay, Gabrielle Brochu, who provided logistics in fieldwork and laboratory. We thank Pierre-André Bordeleau, who provided geomatic support. We thank Johanne Tremblay, Dany Bouchard and Mathieu Michaud, who provided analytic support in isotopy and sugar quantification. The research was funded by the Québec Ministries (Agriculture, Fisheries and Food; Environment, Fight against Climate Change, Fauna and Parks) and by the Natural Sciences and Engineering Research Council of Canada (Alliance program), through two multidisciplinary research initiatives: le pôle d’expertise multidisciplinaire en gestion durable du littoral du Lac Saint-Pierre (2019-2023) and Carbon cycling in Québec’s wetlands (Carbonique, 2024-2029).

