## Supplementary Material for "Lake floodplains as sinks for stable soil organic carbon: is the carbon plant- or microbe-derived, and why does permanent land use matter?"

**Table S1** The mean δ^13^C of current vegetation in each land use

| Land use | Dominant vegetation | Current plant δ^13^C (‰) |
| --- | --- | --- |
| Conventional croplands | Maize (C4) and soybean (C3) | -20.34* |
| Improved croplands | Maize (C4) and soybean (C3) | -20.34* |
| Temporary meadows | Herbaceous (C3) | -23.44 |
| Permanent meadows | Herbaceous (C3) | -27.09 |
| Marshes | Herbaceous (C3) | -29.70 |
| Forested swamps | Tree species (C3) | -30.83 |

*The plant δ^13^C in both croplands was obtained from the weighted average of the δ^13^C values of maize and soybean using their aboveground biomass.

T**able S2** Effects of land use on cumulative SOC stock along the soil profile of 0-100 cm depth

| **Soil depth** | ***P* (SOC concentration)** | ***P* (BD)** | ***P* (soil δ^13^C)** | ***P* (SOC stock)** | ***P* (Cumulative SOC stock*)** |
| --- | --- | --- | --- | --- | --- |
| 0-5 | **<0.001** | **<0.001** | **<0.001** | **<0.01** | **<0.01** |
| 5-10 | **<0.01** | **<0.001** | **<0.001** | 0.07 | **0.02** |
| 10-15 | 0.19 | **<0.001** | **<0.001** | 0.30 | 0.06 |
| 15-20 | 0.58 | **<0.01** | **<0.001** | 0.28 | 0.13 |
| 20-25 | 0.46 | 0.49 | **<0.01** | 0.15 | 0.26 |
| 25-30 | 0.57 | 0.61 | 0.07 | 0.35 | 0.31 |
| 30-35 | 0.16 | 0.51 | 0.20 | 0.12 | 0.28 |
| 35-40 | 0.44 | 0.30 | 0.13 | 0.37 | 0.28 |
| 40-50 | 0.13 | 0.06 | 0.11 | 0.20 | 0.39 |
| 50-60 | 0.50 | 0.37 | **0.02** | 0.87 | 0.38 |
| 60-80 | 0.91 | 0.56 | **0.02** | 0.93 | 0.35 |
| 80-100 | 0.13 | 0.46 | 0.12 | 0.06 | 0.05 |

*Cumulative SOC stock at a given depth is the sum of SOC stocks at that depth and those above it. Significant effects (*P* < 0.05) are highlighted in bold.

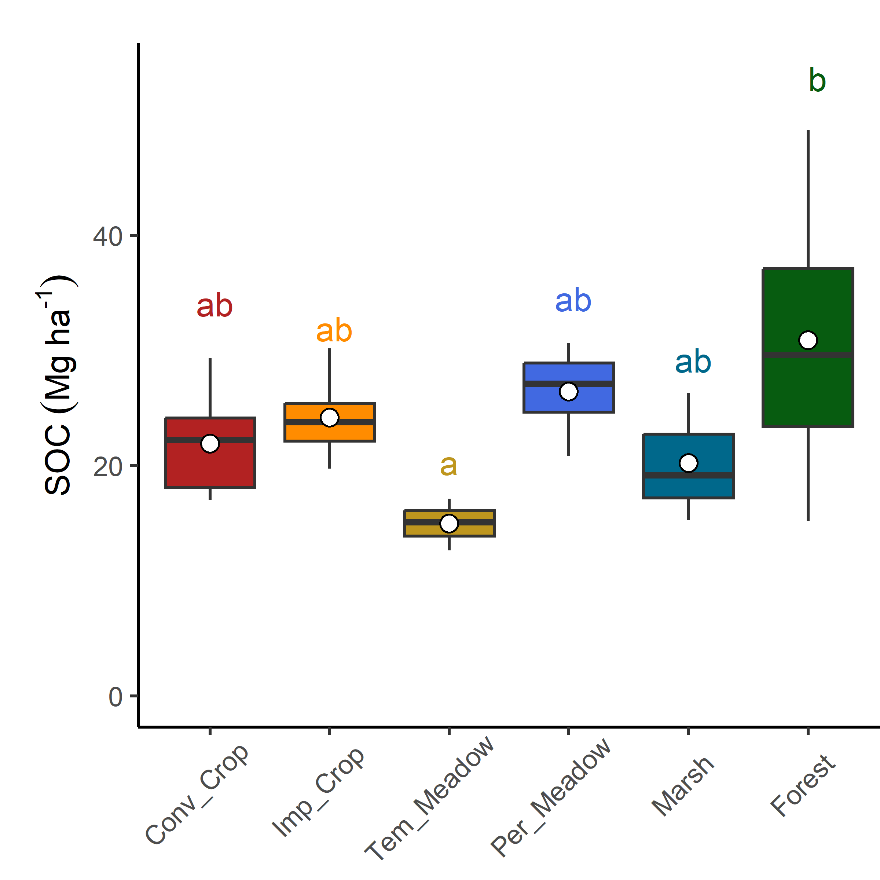
**Fig. S1.** Differences in cumulative SOC stock (0-10 cm) between land uses

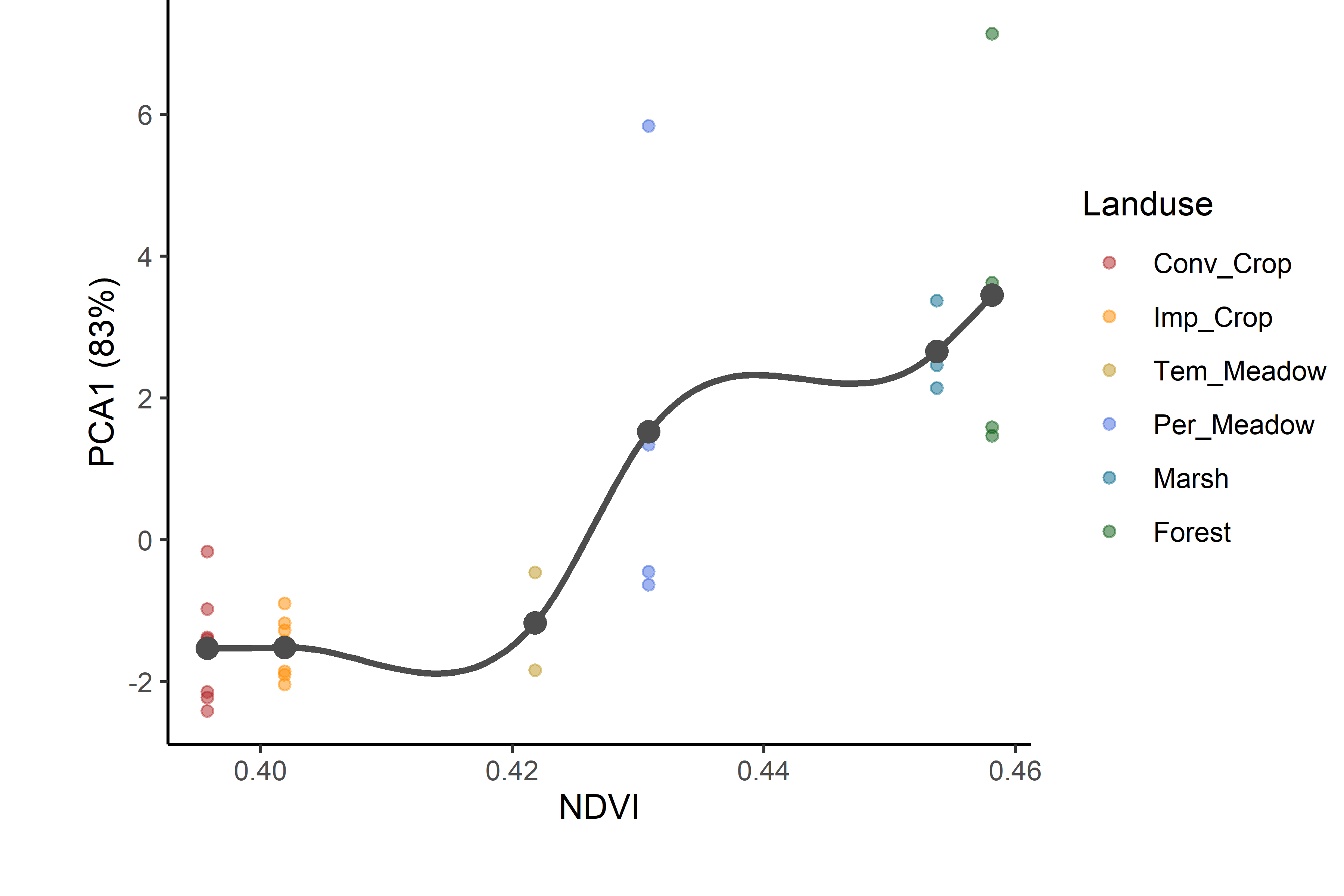
**Fig. S2.** Soil C gain along the anthropogenic disturbance gradient at 0–10 cm depth using Normalized Difference Vegetation Index (NDVI) (b). PCA1 was obtained from principal component analysis (PCA) of soil C variables (see Fig. 6 for details). Conv: conventional; Imp: improved; Per: permanent; Tem: temporary.
